# Conformation and membrane interaction studies of the potent antimicrobial and anticancer peptide palustrin-Ca

**DOI:** 10.1101/2021.07.22.453375

**Authors:** Patrick B. Timmons, Chandralal M. Hewage

**Affiliations:** UCD School of Biomolecular and Biomedical Science, UCD Centre for Synthesis and Chemical Biology, UCD Conway Institute, University College Dublin, Dublin 4, Ireland

## Abstract

Palustrin-Ca (GFLDIIKDTGKEFAVKILNNLKCKLAGGCPP) is a host defense peptide with potent antimicrobial and anticancer activities, first isolated from the skin of the American bullfrog *Lithobates catesbeianus*. The peptide is 31 amino acid residues long, cationic and amphipathic. Two-dimensional NMR spectroscopy was employed to characterise its three-dimensional structure in a 50/50% water/2,2,2-trifluoroethanol-*d_3_* mixture. The structure is defined by an *α*-helix that spans between Ile^6^-Ala^26^, and a cyclic disulphide bridged domain at the C-terminal end of the peptide sequence, between residues 23 and 29.

A molecular dynamics simulation was employed to model the peptide’s interactions with sodium dodecyl sulphate micelles, a widely used bacterial membrane-mimicking environment. Throughout the simulation, the peptide was found to maintain its *α*-helical conformation between residues Ile^6^-Ala^26^, while adopting a position parallel to the surface to micelle, which is energetically-favourable due to many hydrophobic and electrostatic contacts with the micelle.

## 1 Introduction

Host defence peptides (HDPs) are naturally occurring molecules that are secreted as part of the non-specific response of the innate immune system^[1]^. These molecules contribute to the immune system’s antimicrobial response either by exerting direct antimicrobial activity, or by altering the host’s immune response. HDPs are particularly prevalent in amphibians, with approximately one third of all known HDPs having been originally isolated from amphibians. Nonetheless, HDPs are a highly conserved part of the immune response, and have been found in all species of organisms, from humans to bacteria^[2–5]^.

As the number of documented cases of antimicrobial resistance to conventional small-molecule drugs continues to rise, interest in HDPs, which can exert their antimicrobial activity against both Gram-positive bacteria and Gram-negative bacteria without the development of resistance, continues to grow^[6]^. HDPs possess a number of additional advantages. They are more efficacious, selective and specific than small molecules, and are degraded to naturally occurring amino acids, which reduces the danger of unfavourable drug-drug interactions. HDPs are suitable for use both in combination with conventional drugs, and as standalone substitutes. The biological activities of HDPs, however, are not limited to bacteria. HDPs have also been demonstrated to possess antifungal, antiparasitic, antiviral and anticancer activities^[7]^. Indeed, palustrin-Ca (GFLDIIKDTGKEFAVKILNNLKCKLAG-GCPP) is reported to be the most potent known anticancer peptide, with an IC_50_ of 0.951 *μg/ml* against human gastric cancer SGC-7901^[8]^.

Typically, HDPs are short, with a primary sequence length between 10-50 amino acid residues. Usually, HDPs are unstructured in aqueous solution, and only adopt a defined secondary structure upon contact with a biological membrane, or in a hydrophobic environment. The induced structures are predominantly *α*-helical in nature, although other identified structures include *β*-sheet, mixed *α*-*β* and extended structures. A common feature of HDP structures are that they are typically amphipathic in nature, with a clear separation between the peptide’s hydrophobic and polar or charged residues. Overall, HDP’s are usually positively charged, although a number of negatively charged peptides have been isolated as well^[9]^. Indeed, increased positive charge, up to a certain extent, has been associated with greater antimicrobial activity^[10,11]^. Similarly, increased peptide hydrophobicity is associated with peptide hemolytic activity, and thereby represents another key feature which can be changed to modulate peptide selectivity^[12,13]^.

While HDPs are typically positively charged, anionic molecules such as lipopolysaccharides (LPS), phospholipids and teichoic acids are prevalent in prokaryotic cell membranes, which enables attractive electrostatic interactions between HDPs and their target bacterial cell membranes^[14,15]^. Similarly to prokaryotic membranes, the cell membranes of cancer cells are enriched in negative charge, unlike non-cancerous cell membranes which are neutral, which allows anticancer peptides to be selectively electrostatically attracted to cancer cells.

HDPs usually exert their bactericidal activity via disruption of the bacterial cell membrane, for which three main mechanisms of action have been described: the toroidal pore model, barrel-stave model and carpet model. All three membranolytic mechanisms of action are relatively non-specific, which is the reason why few cases of bacterial resistance to HDPs have been described^[16,17]^. The initial step in all three mechanisms is the HDP binding of the target membrane, initially facilitated via the aforementioned electrostatic attraction, then via hydrophobic attraction. Eventually, a threshold peptide concentration is reached, which may be determined by the organism’s capacity to repair its damaged cell membranes^[18]^, whereupon the mechanism proceeds via the pore-forming toroidal pore or barrel-stave models, or the non-pore-forming carpet model. The latter requires that the bilayer curvature is disrupted, thereby disintegrating the bacterial membrane. Under the toroidal pore model, the bacterial membrane is bent from the outer leaflet inwards, resulting in the formation of a transmembrane toroidal pore lined by the peptides and the membrane’s lipid head groups^[19,20]^. The barrel-stave model, meanwhile, involves the peptides forming a transmembrane pore; its lumen is lined by the peptides’ hydrophobic residues, and its exterior is composed of hydrophilic residues, which are in contact with the membrane’s lipid headgroups. All the models of HDP activity culminate in the dissipation of the cross-membrane electrical potential and the leakage of cellular components.

Our laboratory has conducted structural studies on various amphibian HDPs, including ranatuerin-2CSa^[21]^, XT-7^[22]^, alyteserin-1c^[23]^, brevinin-1BYa^[24]^ and its analogues^[25]^, maximin 3^[26]^ and maximin 1^[27]^. Palustrin-Ca (GFLDIIKDTGKEFAVKILNNLKCKLAGGCPP) is a 31 amino acid residue HDP first isolated from the skin of the American bullfrog *Lithobates catesbeianus* ^[8]^. In common with other HDPs isolated from frogs belonging to the Ranidae family, it contains a cyclic disulphide bridged domain at the C-terminal end of the peptide sequence, between residues 23 and 29. Palustrin-Ca is biologically interesting, as it non-hemolytic, and exhibits potent broad-spectrum antibacterial activity, and as previously mentioned, very potent anticancer activity, with an IC_50_ of 0.951 *μg/ml* against human gastric cancer SGC-7901^[8]^. In this work, palustrin-Ca’s conformation and structural properties are elucidated using NMR spectroscopy, with an ensemble of model structures being determined. Additionally, the peptide’s interactions with a membrane-mimetic are simulated using an atomistic molecular dynamics simulation.

## 2 Materials and methods

### 2.1 Materials and NMR sample preparation

The palustrin-Ca peptide (MW=3304 g mol^-1^, purity *>*95%) was purchased from ProteoGenix (Paris). 4.5 g of the peptide was dissolved in 0.6 mL of a 50% (v/v) TFE-*d_3_*/H_2_O solution, resulting in a final peptide concentration of 2.27 mM. 3-trimethylsilyl propionic acid (TSP) and 2,2,2-trifluoroethanol (TFE-*d_3_*) of analytical grade were obtained from Sigma-Aldrich (Ireland).

### 2.2 NMR spectroscopy

A Bruker Avance 600 NMR spectrometer with a 5mm inverse probe head at a ^1^H resonance frequency of 600.13 MHz was used to perform a number of NMR experiments at a temperature of 298 K. One-dimensional proton, 2D phase-sensitive total correlation spectroscopy (TOCSY)^[28]^, nuclear Overhauser effect spectroscopy (NOESY)^[29]^ and natural abundance ^1^H-^13^C and ^1^H-^15^N heteronuclear single quantum coherence spectroscopy (^1^H-^13^C-HSQC) and ^1^H-^15^N-HSQC^[30]^ spectra were acquired, with relaxation delays of 2.5 s, 2.0 s, 1.5 s, 1.0 s and 2.0 s, respectively, and acquisition times of 3.4 s, 340 ms, 280 ms, 170 ms and 100 ms, respectively. Mixing times of 60 ms and 200 ms were used for the TOCSY and NOESY, respectively.

The ^1^H spectral widths were 6.0 kHz, 7.2 kHz, 6.0 kHz and 9.6 kHz, and the spectra were acquired with 4, 16, 32 and 64 transients for each of the 1024, 2048, 256 and 128 t1 increments for the TOCSY, NOESY, ^1^H-^13^C-HSQC and ^1^H-^15^N-HSQC, respectively. The ^13^C spectral width was 21.1 kHz and the ^15^N spectral width was 10.9 kHz. All two-dimensional spectra were processed with the Bruker TopSpin program, version 4.0.6 (Bruker BioSpin, Germany), using the sine squared window function, and the ^1^H signal of TSP was used as the chemical shift reference.

### 2.3 Structure calculation

The NMRFAM-SPARKY, version 3.131 ^[31]^ was used to analyse the acquired NMR spectra and integrate the NOESY cross-peaks. The integrated peak volumes were exported, and the nuclear Overhauser effect (nOe) cross-peak intensities were calibrated using the CALIBA^[32]^ program, yielding a set of upper distance restraints. Protons that could not be stereospecifically assigned were treated as pseudoatoms. The set of obtained distance restraints was used as input to CYANA^[33]^, excluding those that represent fixed distances. A force constant of 1 kJ mol^-1^ Å^-2^ was used to weight the distance restraints, from which one hundred structures were generated by CYANA^[34]^, and submitted to 20,000 steps of simulated annealing and 20,000 of conjugate gradient minimization. The 20 structures with the lowest target function values were selected, and subjected to an additional 2,000 steps of conjugate gradient energy minimization with restrained backbone atoms, using the CHARMM22 force field^[35]^ in NAMD, version 2.12^[36]^. VMD (visual molecular dynamics), version 1.9.3^[37]^, was used to analyse the final ensemble of 20 structures. The geometry, stereochemical quality and structural statistics of the final ensemble of structures was assessed and validated using PROCHECK^[38]^ and the wwPDB web service^[39]^.

### 2.4 Molecular dynamics simulation

Structural model coordinates of an SDS micelle were obtained from Jakobtorweihen et al.^[40]^, and used in the construction of an SDS micelle-peptide system, which was constructed by aligning the centres of mass of the micelle and the medoid palustrin-Ca structure with VMD. To ensure that the system’s charge remained neutral, chloride ions were added as counter-ions, and finally, TIP3P water was used to solvate the system.

The system was energy minimized and equilibrated using the CHARMM22 all-atom forcefield^[35,41]^ within NAMD version 2.12 ^[36]^. All calculations were conducted within the NPT ensemble, using the Langevin piston Nose-Hoover method^[42,43]^ and periodic boundary conditions. Long-range non-bonded interactions were calculated up to a switching distance of 8.5 Å, beyond which a smooth switching function truncated the energy to a cut-off of 11 Å. The PME method^[44]^ was used to calculate long-range electrostatic interactions at each time step. The non-bonded interaction list was also updated every step. All bonds to hydrogen atoms were constrained using the SHAKE algorithm^[45]^. A 2 fs timestep was employed.

The solvated peptide-micelle system was minimized for 2,000 conjugate gradient steps with peptide backbone atoms fixed, and a further 2,000 steps with its *α*-carbon atoms restrained. The system was then heated to a temperature of 310 K over 6,000 steps using Langevin dynamics, with *α*-carbon remaining restrained. The system volume was equilibrated with the Langevin piston at 1 atm over 24,000 steps, with a further 24,000 steps without restraints. Finally, the system was simulated for 57.4 ns.

The radial distribution function *g*(*r*) was used for analysing the final XX ns of simulation data. The *measure gofr* command within VMD was used to calculate the *g*(*r*) function between each residue’s carbon atoms and the SDS micelle’s non-sulfate atoms. The default values 0.1 and 25.0 Å were used for *δr* and max *r*, respectively.

## 3 Results and Discussion

### 3.1 Conformational analysis by NMR

Two-dimensional NMR spectroscopy was used to determine the structure of palustrin-Ca in a 50:50 TFE-*d_3_*/water solvent mixture, which is known to promote the formation of secondary structural elements observed in biological membranes by enhancing the peptide’s *α*-helical character^[46]^. The collected spectra were well-resolved, with well-dispersed peaks that facilitated unambiguous resonance assignment. Individual residue spin systems were identified with the aid of TOCSY, ^1^H-^13^C HSQC and ^1^H-^15^N HSQC spectra, while sequence-specific resonance assignment was conducted using the HN-HN and HN-H*α* regions of the NOESY spectrum. The first sequence-specific resonance assignments were made for the unique residues Thr^9^, Glu^12^ and Val^15^, with the remainder being assigned through the use of a backbone-walk, and investigation of HN-H*α* region peaks, where necessary. The Gly^1^ amide proton chemical shift could not be determined due to chemical exchange. The presence of a disulfide bridge was confirmed by observation of clear long-range *β_i_*-*β_i+6_* nOe peaks between residues Cys^23^-Cys^29^.

**Figure 1:**
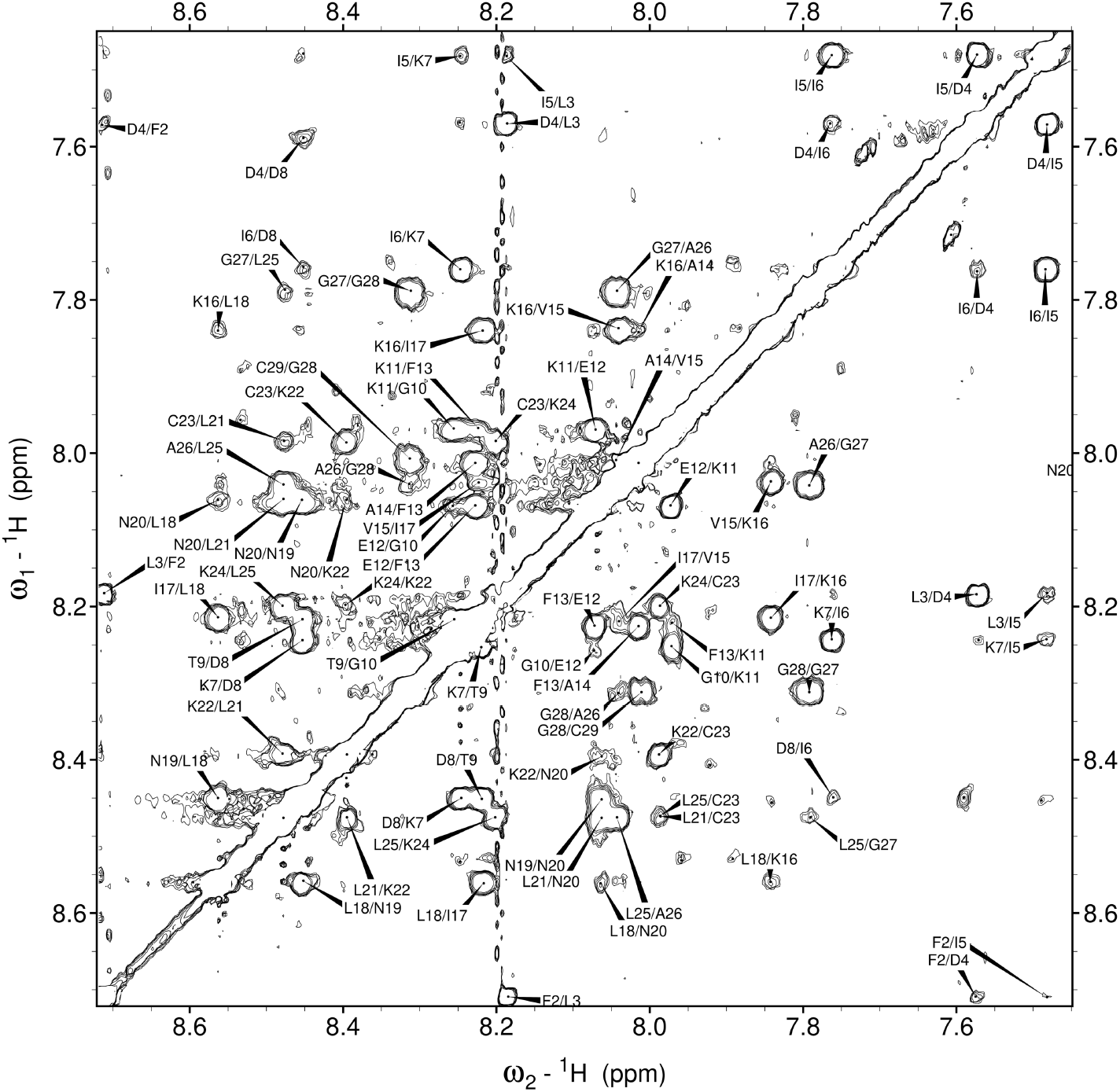
Amide region of the 200 ms NOESY spectrum of palustrin-Ca in 50% TFE-*d_3_*-H_2_O mixed solvent system with d_NN_(*i,i+1*) connectivities labelled.

**Figure 2:**
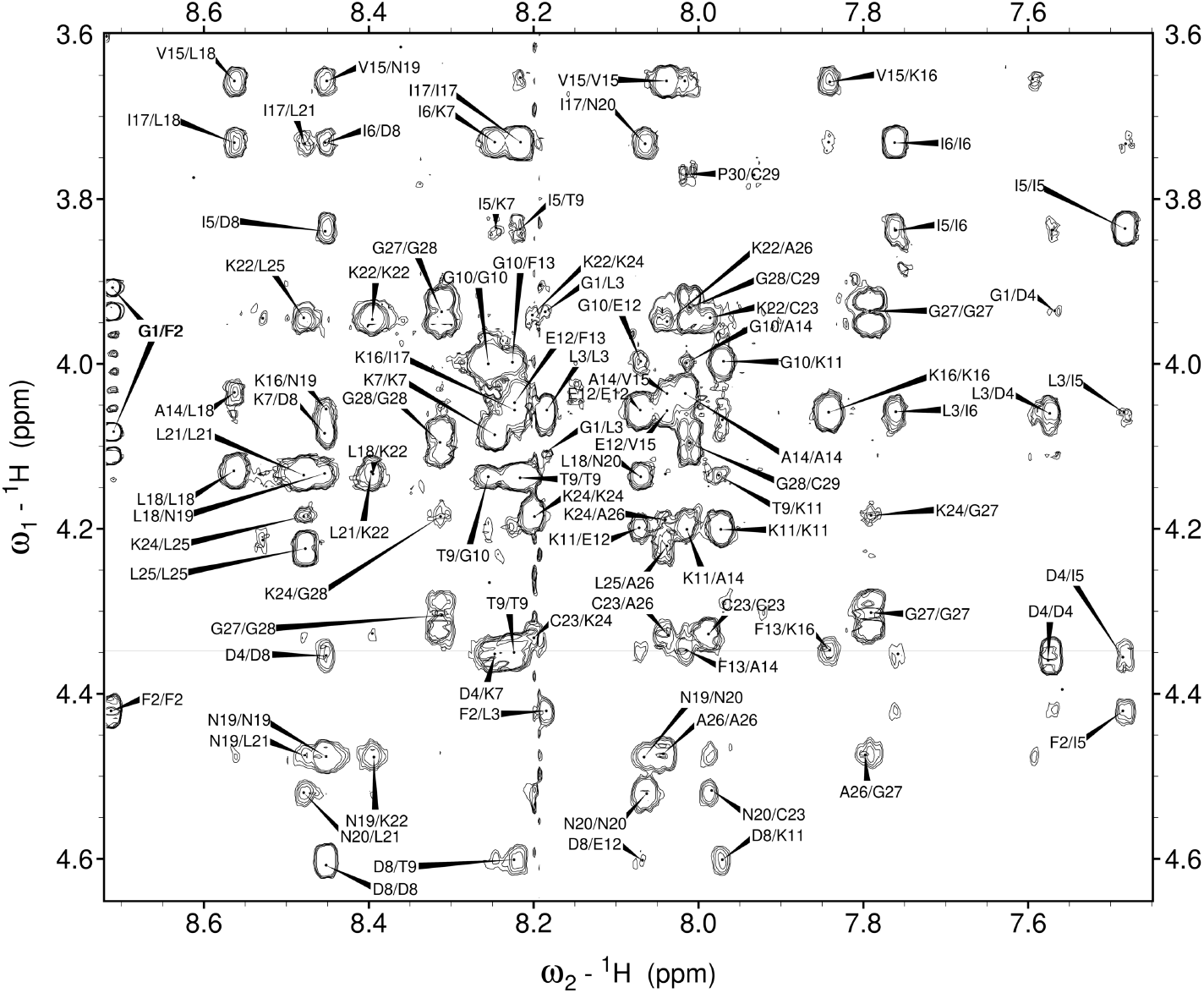
Fingerprint region of the 200 ms NOESY spectrum of palustrin-Ca in 50% TFE-*d_3_*-H_2_O mixed solvent system with backbone connectivities labelled.

All the ^1^H chemical shifts identified for palustrin-Ca are detailed in Table 1. Likewise, all of palustrin-Ca’s intermolecular nOe connectivities are summarised in Figure 3, where the nOe intensity is proportional to the thickness of each line. Multiple d_*α*N_(i, i+3), d*_αβ_*(i, i+3), d_*α*N_(i, i+4) and d*_αβ_*(i, i+4) were identified, which are indicative of an *α*-helical secondary structure.

**Table 1:** ^1^H chemical shifts (ppm) identified for every residue of palustrin-Ca in 50% TFE-*d_3_*/H_2_O

| Amino acid | NH | Hα | Hβ | Other protons |
| --- | --- | --- | --- | --- |
| Gly<sup>1</sup> |  | 4.101, 3.923 |  |  |
| Phe<sup>2</sup> | 8.711 | 4.421 | 3.240, 3.075 | $\delta$ 7.235; $\epsilon$ 7.321 |
| Leu<sup>3</sup> | 8.185 | 4.059 | 1.683, 1.569 | $\gamma$ 1.634; $\delta$ 0.972, 0.920 |
| Asp<sup>4</sup> | 7.573 | 4.356 | 2.880, 2.777 |  |
| Ile<sup>5</sup> | 7.482 | 3.836 | 2.064 | $\gamma$ 1.251, 0.917; $\delta$ 0.922; |
| Ile<sup>6</sup> | 7.761 | 3.731 | 1.970 | γ1 1.482, 1.106; γ2 0.893; $\delta$ 0.808 |
| Lys<sup>7</sup> | 8.246 | 4.085 | 1.939 | $\gamma$ 1.666, 1.486; $\delta$ 1.733, $\epsilon$ 2.980 |
| Asp<sup>8</sup> | 8.452 | 4.603 | 3.075, 2.941 |  |
| Thr<sup>9</sup> | 8.219 | 4.351 | 4.136 | $\gamma$ 1.293 |
| Gly<sup>10</sup> | 8.254 | 3.998 |  |  |
| Lys<sup>11</sup> | 7.971 | 4.198 | 2.005 | $\gamma$ 1.655, 1.525; $\delta$ 1.764; $\epsilon$ 3.021, 3.005 |
| Glu<sup>12</sup> | 8.070 | 4.056 | 2.288, 2.172 | $\gamma$ 2.606, 2.457 |
| Phe<sup>13</sup> | 8.226 | 4.348 | 3.272 | $\delta$ 7.277, 7.291 |
| Ala<sup>14</sup> | 8.014 | 4.035 | 1.587 |  |
| Val<sup>15</sup> | 8.039 | 3.656 | 2.222 | $\gamma$ 1.125, 0.989 |
| Lys<sup>16</sup> | 7.841 | 4.059 | 2.058, 1.964 | $\gamma$ 1.478, 1.449; $\delta$ 1.716, 1.676; $\epsilon$ 3.007, 2.994 |
| Ile<sup>17</sup> | 8.217 | 3.732 | 1.979 | γ1 1.499, 1.071; γ2 0.888; $\delta$ 0.725 |
| Leu<sup>18</sup> | 8.561 | 4.132 | 1.925 | $\gamma$ 1.583; $\delta$ 0.922 |
| Asn<sup>19</sup> | 8.453 | 4.477 | 2.972, 2.771 | $\delta$ 6.588 |
| Asn<sup>20</sup> | 8.063 | 4.521 | 3.051, 2.831 | $\delta$ 6.789 |
| Leu<sup>21</sup> | 8.477 | 4.134 | 1.919, 1.844 | $\gamma$ 1.781; $\delta$ 0.948, 0.914 |
| Lys<sup>22</sup> | 8.395 | 3.944 | 2.003 | $\gamma$ 1.455; $\delta$ 1.720, 1.673; $\epsilon$ 2.978 |
| Cys<sup>23</sup> | 7.987 | 4.328 | 3.472, 3.185 |  |
| Lys<sup>24</sup> | 8.200 | 4.185 | 2.091, 1.988 | $\gamma$ 1.645, 1.517; $\delta$ 1.714 |
| Leu<sup>25</sup> | 8.476 | 4.222 | 1.892 | $\gamma$ 1.544; $\delta$ 0.906 |
| Ala<sup>26</sup> | 8.040 | 4.471 | 1.546 |  |
| Gly<sup>27</sup> | 7.791 | 4.304, 3.936 |  |  |
| Gly<sup>28</sup> | 8.312 | 4.091, 3.936 |  |  |
| Cys<sup>29</sup> | 8.012 | 4.970 | 3.377, 2.951 |  |
| Pro<sup>30</sup> | | 4.662 | 2.324, 2.064 | $\gamma$ 2.006; $\delta$ 3.768, 3.661 |
| Pro<sup>31</sup> | | 4.391 | 2.313, 2.058 | $\gamma$ 2.010; $\delta$ 3.766, 3.622 |

**Figure 3:**
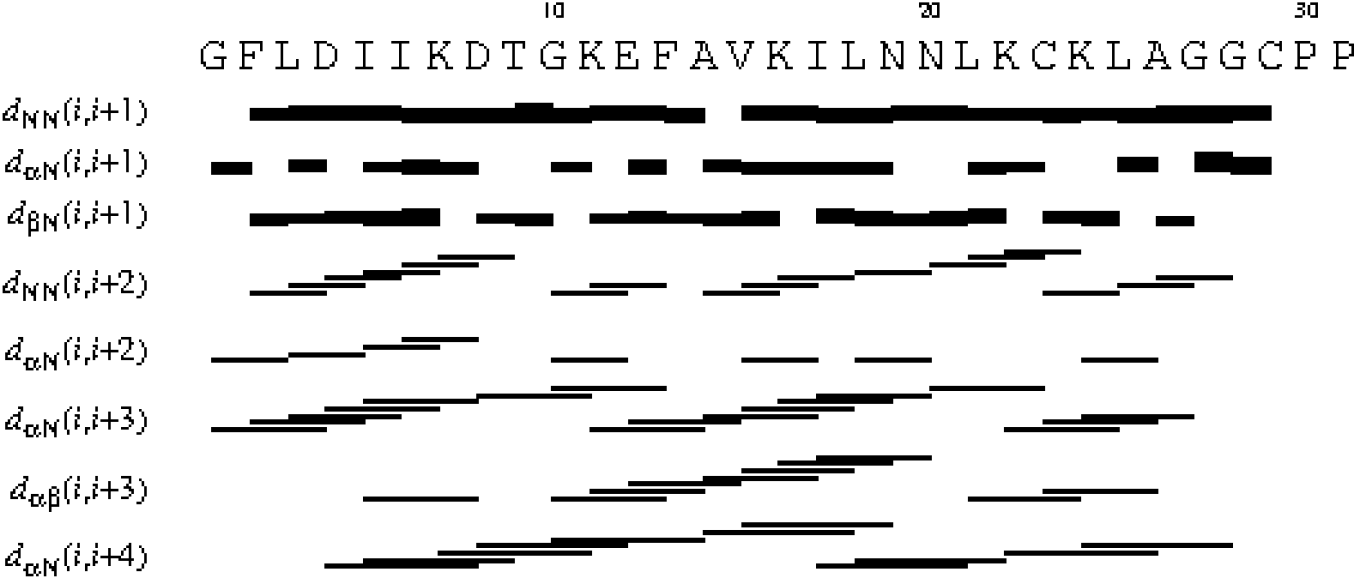
Short- and medium-range connectivities for palustrin-Ca in 50% TFE-*d_3_*-H_2_O.

### 3.2 Chemical shift analysis

Amide proton chemical shift deviations, calculated as the difference between the observed chemical shift and the random coil chemical shift (Δ*δ = δ_obs_ δ_rc_*), are connected with the hydrogen bond length^[47]^. Shorter hydrogen bond lengths are associated with higher Δ*δ* values, and vice versa. Calculated amide proton chemical shift deviations are illustrated in Figure 4a; the corresponding hydrogen bond lengths are given in Figure 4b. The amide proton chemical shift deviations exhibit a periodicity of 3-5 from residue 8 on, with greater frequency in the C-terminal part of the sequence. The greatest values are found for the Phe^2^, Asp^8^, Phe^13^, Leu^18^, Leu^21^, Leu^25^ and Gly^28^, while the lowest values are observed for Asp^4^, Lys^11^, Lys^16^, Asn^20^, Cys^23^ and Gly^27^. As expected, the greatest values are observed for hydrophobic residues, with the exception of the Asp^8^ and Gly^28^ residues. Similarly, the lowest values are observed for polar and charged residues.

**Figure 4:**
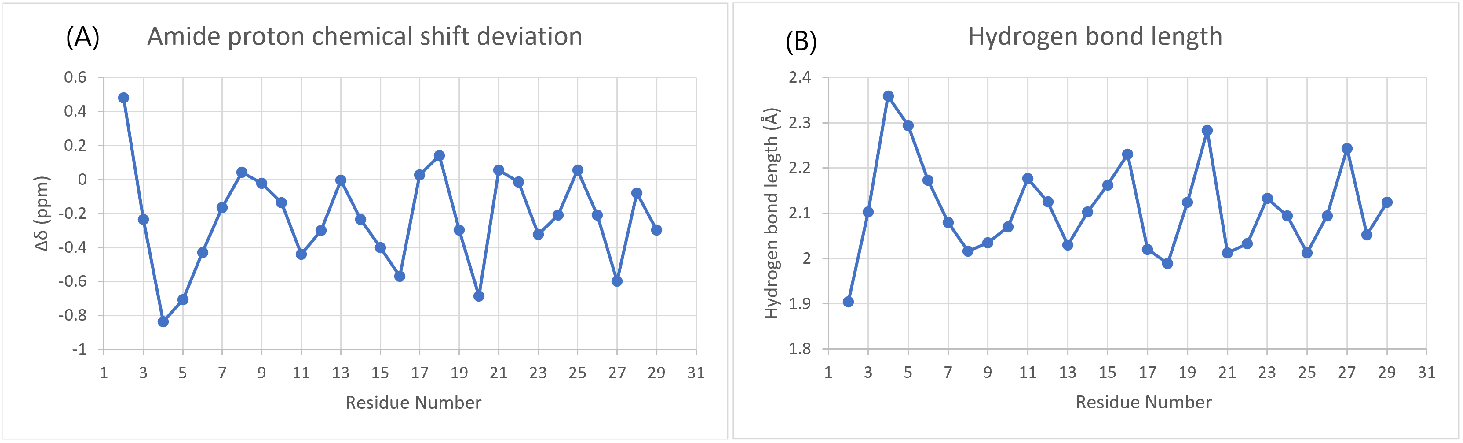
**a** Amide proton chemical shift deviation plot of palustrin-Ca in 50% TFE-*d_3_*-H_2_O. The observed amide proton chemical shifts were compared to the standard random coil chemical shifts (Δ*δ = δ_obs_ − δ_rc_*. **b** Hydrogen bond distance as calculated from the chemical shift deviations 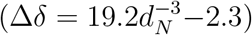, where *d_N_* denotes hydrogen bond length^[47]^.

Similarly, *α*-proton chemical shift deviations were calculated as the difference between the observed chemical shift and the random coil chemical shift (Δ*δ = δ_obs_ δ_rc_*). Four sequential *α*-proton chemical shift deviations at least 0.1 ppm lower than the expected random coil values indicate *α*-helical segments, while three or more deviations at least 0.1 ppm higher are indicative of *β*-strands; no change greater than 0.1 ppm is indicative of coiled regions^[48]^. Figure 5 shows the chemical shift deviations of palustrin-Ca, which shows a clear stretch of downfield shifts between residues 2-8, and again between residues 11-25. While these chemical shift deviations are indicative of *α*-helical structure, the former stretch of downfield shifts may instead be attributed to a turn conformation.

**Figure 5:**
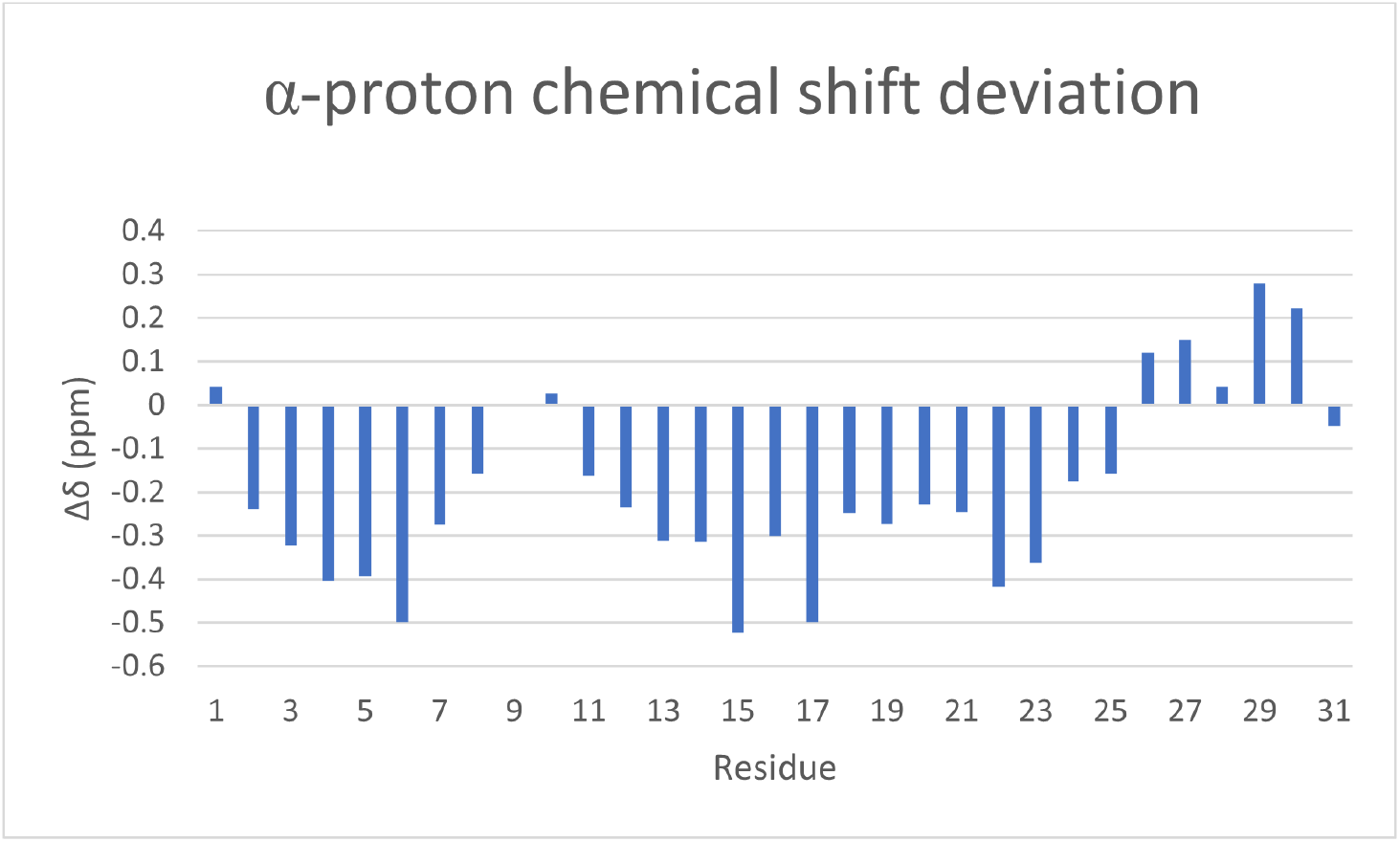
*α*-proton chemical shift deviation plot of palustrin-Ca in 50% TFE-*d_3_*-H_2_O. The observed *α*-proton chemical shifts were compared to the standard random coil chemical shifts (Δ*δ = δ_obs_ − δ_rc_*.

### 3.3 Molecular modelling

The NOESY cross-peaks were integrated, and the resultant volumes were converted into distance restraints. CYANA was employed to generate one hundred structures, of which the twenty possessing the lowest target function values were further energy-minimized. Table 2 summarizes the structural and energetic statistics of the twenty selected models. As depicted in Figure 6 is predominantly *α*-helical. The ensemble’s secondary structural properties were analysed with the aid of STRIDE^[49]^. Most of the ensemble’s structures are display *α*-helical structures between residues Ile^6^-Ala^26^. The terminal segments are predominantly defined as turns, but possess significant coil character as well.

**Table 2:**
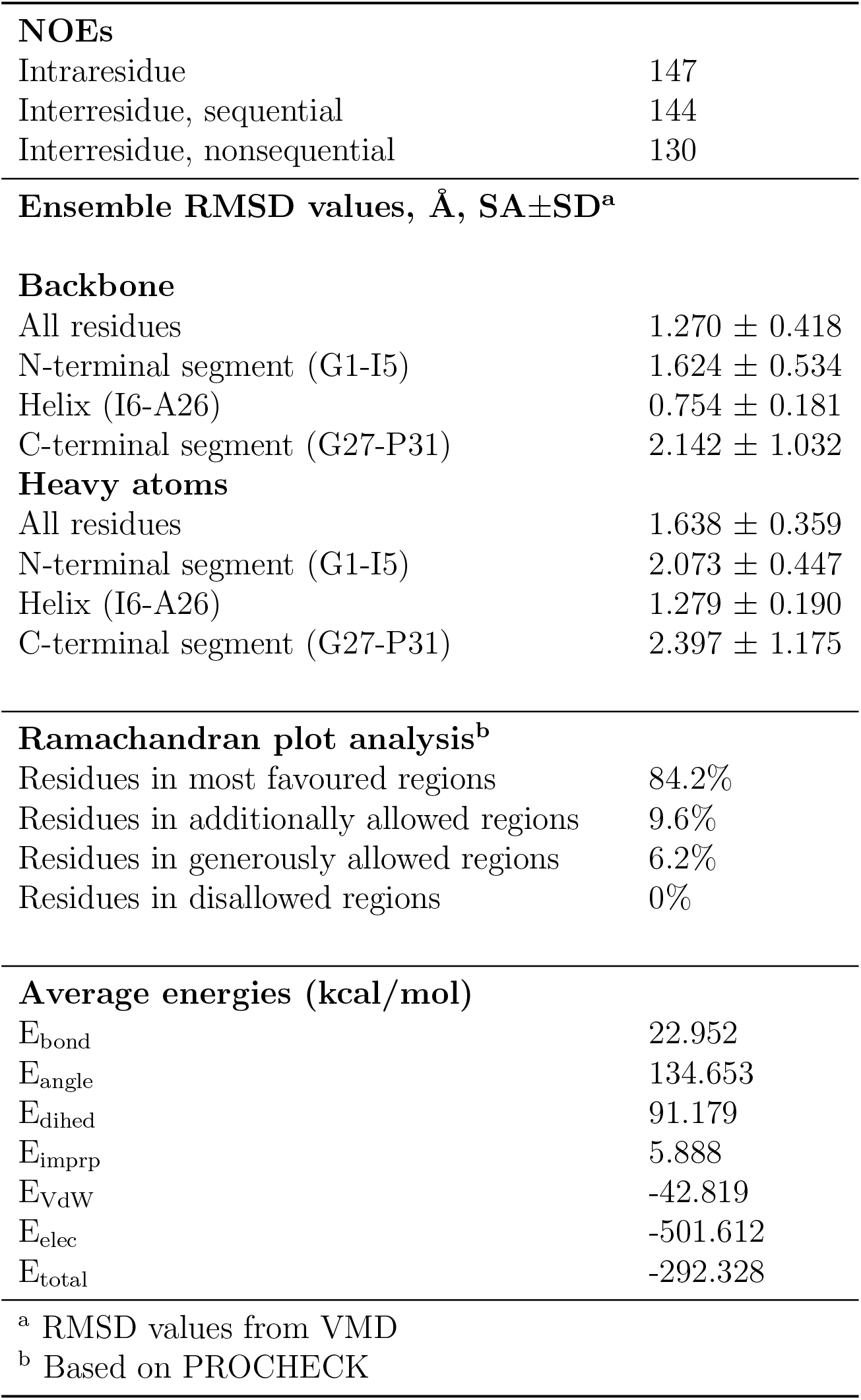
Mean structural statistics of the twenty structural models comprising the palustrin-Ca ensemble. The peptide structure has been deposited in the PDB with deposition code 7P4X. Model 10 is the representative medoid structure.

|  |  |
| --- | --- |
| **NOEs** |  |
| Intraresidue | 147 |
| Interresidue, sequential | 144 |
| Interresidue, nonsequential | 130 |
| **Ensemble RMSD values, Å, SA±SD<sup>a</sup>** |  |
| **Backbone** |  |
| All residues | 1.270 ± 0.418 |
| N-terminal segment (G1-I5) | 1.624 ± 0.534 |
| Helix (I6-A26) | 0.754 ± 0.181 |
| C-terminal segment (G27-P31) | 2.142 ± 1.032 |
| **Heavy atoms** |  |
| All residues | 1.638 ± 0.359 |
| N-terminal segment (G1-I5) | 2.073 ± 0.447 |
| Helix (I6-A26) | 1.279 ± 0.190 |
| C-terminal segment (G27-P31) | 2.397 ± 1.175 |
| **Ramachandran plot analysis<sup>b</sup>** |  |
| Residues in most favoured regions | 84.2% |
| Residues in additionally allowed regions | 9.6% |
| Residues in generously allowed regions | 6.2% |
| Residues in disallowed regions | 0% |
| **Average energies (kcal/mol)** |  |
| E<sub>bond</sub> | 22.952 |
| E<sub>angle</sub> | 134.653 |
| E<sub>dihed</sub> | 91.179 |
| E<sub>imprp</sub> | 5.888 |
| E<sub>VdW</sub> | -42.819 |
| E<sub>elec</sub> | -501.612 |
| E<sub>total</sub> | -292.328 |
| <sup>a</sup> RMSD values from VMD |  |
| <sup>b</sup> Based on PROCHECK |  |

**Figure 6:**
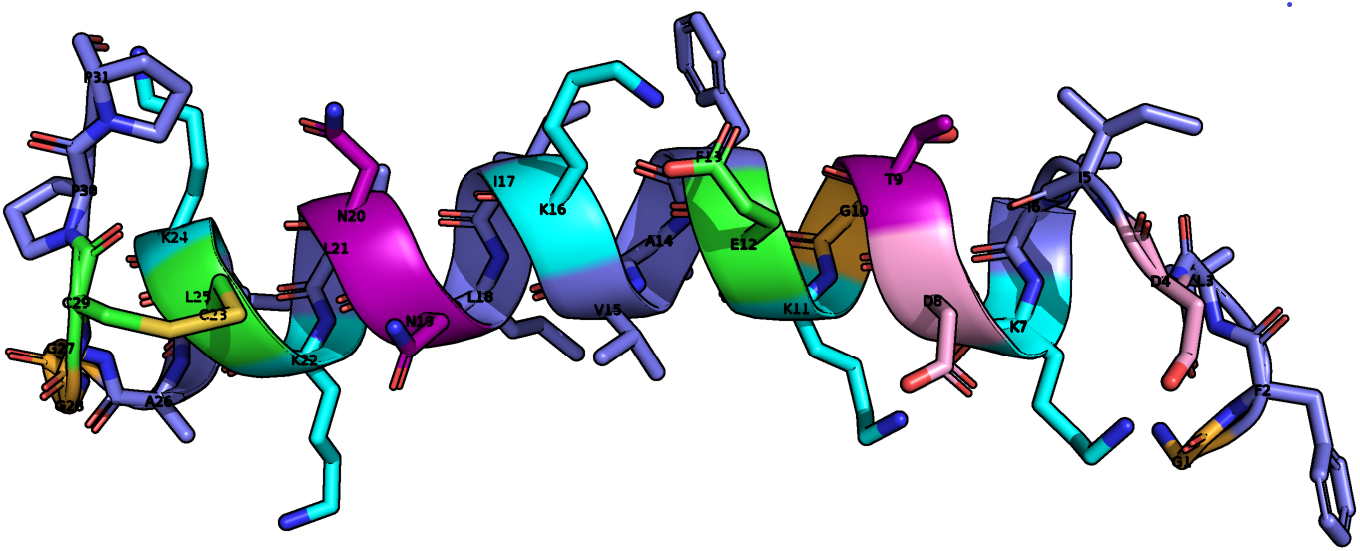
Ribbon representation of the medoid solution structure of palustrin-Ca in 50% TFE-d_3_-H_2_O. The amphipathic helical motif is observed. Positively charged residues are coloured in pink, negatively charged residues coloured in blue, polar residues coloured in purple, glycine residues coloured in orange, cysteine residues are coloured in green, hydrophobic residues coloured in slate.

### 3.4 Molecular dynamics

A molecular dynamics simulation of palustrin Ca’s medoid structure in an SDS micelle was conducted to characterise its behaviour in the prokaryotic membrane-mimetic environment. In the course of the simulation, the peptide translocated from its initial position aligned with the micelle’s centre of mass, to the micelle’s surface-water boundary, where it adopted a position parallel to the micelle surface (Figure 7). The peptide’s amphiphilicity is clearly apparent; its hydrophobic residues remain closely associated with the micellar interior, while the hydrophilic residues are in contact with the aqueous solvent and the micelle’s anionic sulfate headgroups. This configuration is quite energetically favourable.

**Figure 7:**
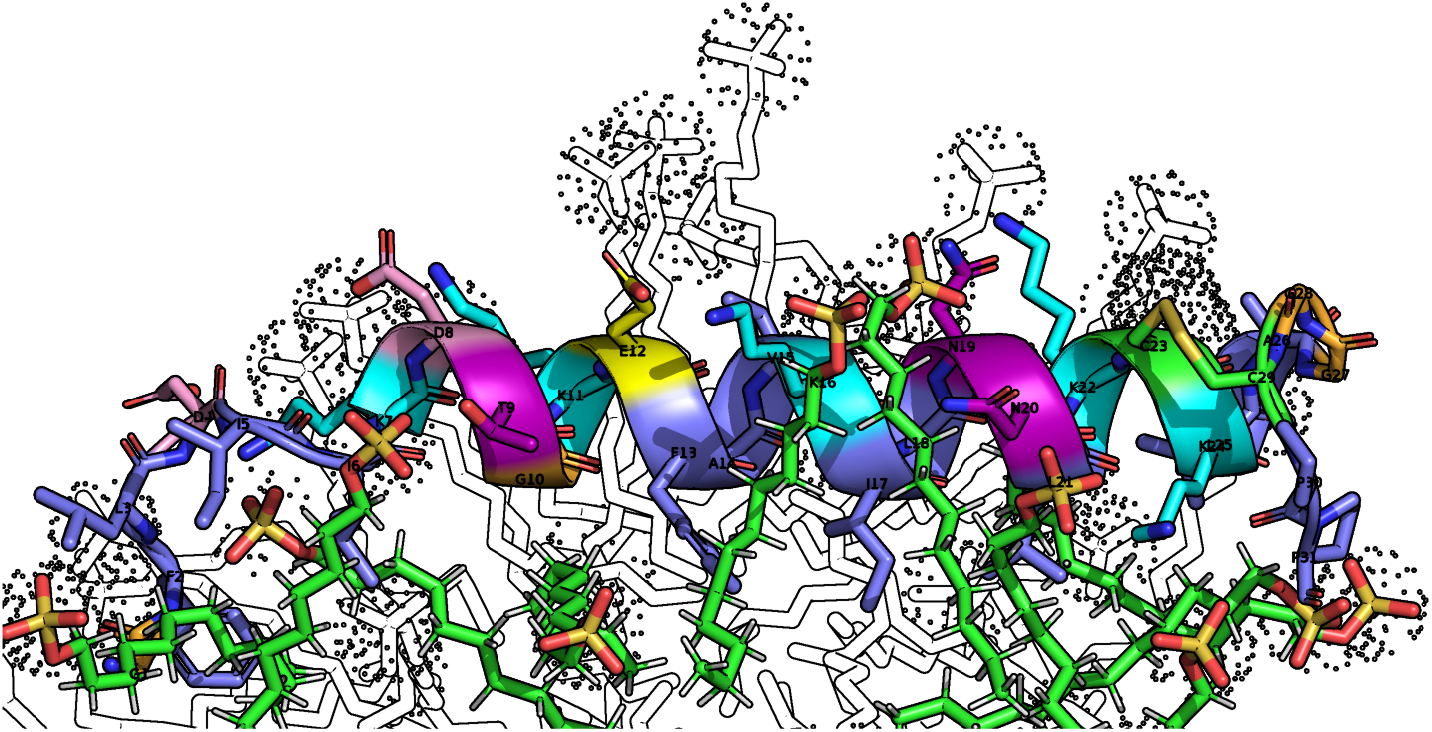
Palustrin-Ca positioned at the SDS micelle-water boundary. Lysine residue side chains are shown interacting with the negatively charged sulfate headgroups. Positively charged residues are coloured in pink, negatively charged residues coloured in blue, polar residues coloured in purple, glycine residues coloured in orange, cysteine residues are coloured in green, hydrophobic residues coloured in slate.

The peptide’s interactions with the micelle’s hydrophobic core were quantified using the radial distribution function *g(r)*, which was calculated between the peptide’s carbon atoms and the micelle’s aliphatic chain, and plotted against the radius *r*, as shown in Figure 8.

**Figure 8:**
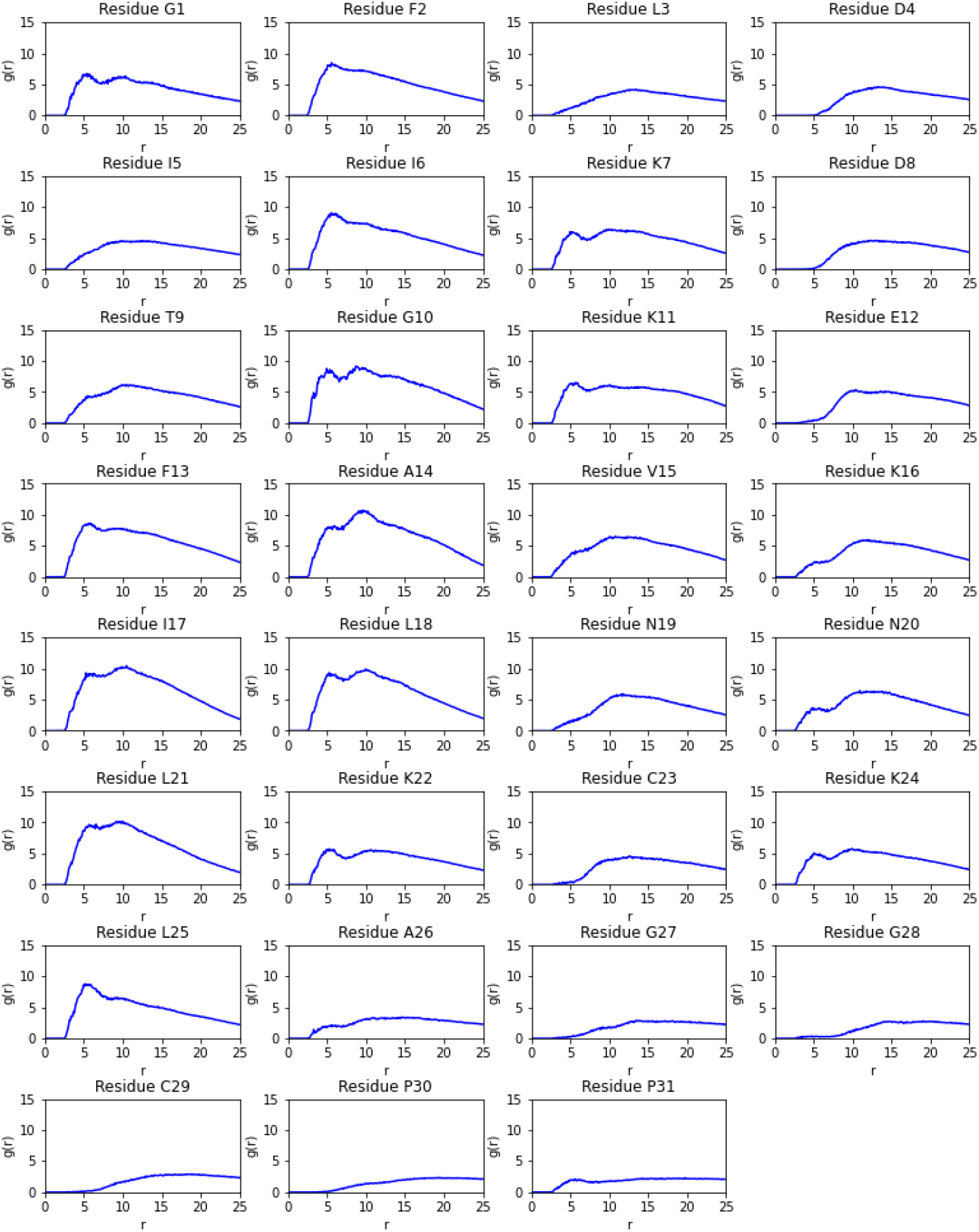
Radial distribution function (RDF) *g(r)* plotted between palustrin-Ca’s residues’ carbon atoms and the sodium dodecyl sulfate (SDS) micelle aliphatic chain.

The radial distribution function plots illustrate clearly that the peptide’s hydrophobic residues maintain a close association with the micelle, especially in the structured *α*-helical segment. Interestingly, the N-terminal segment of the peptide is also found to maintain close association with the micelle, especially through the Gly^1^, Phe^2^ and Ile^6^ residues, despite the first five residues not forming part of a defined secondary structure. This is in contrast to the C-terminal segment, where beginning with Ala^26^ onwards the peptide is only weakly associated with the micelle.

The peptide, which was experimentally determined to be *α*-helical Ile^6^ and Ala^26^, maintains this structure throughout the duration of the simulation. Despite the presence of a glycine residue at position 10, the *α*-helix remains relatively rigid, not exhibiting any significant flexibility during the simulation. This suggests that the glycine residue occurs too far from the helical segment’s middle to impart flexibility. Although studies have previously demonstrated that a rigid *α*-helical structure can potentially result in an increased hemolytic activity^[50]^, this does not appear to apply in this case, which may be explained by the relatively flexible N-terminal and C-terminal segments, which combined account for 10 of the sequences 31 residues.

## 4 Conclusion

A number of biologically active peptides have been isolated from the American bullfrog *Lithobates catesbeianus*. The isolated peptides are diverse, and not limited to a single peptide family; palustrin-Ca is the only palustrin peptide isolated from this amphibian. Palustrin-Ca can be considered to be a member of the palustrin-2 family of peptides, sharing an average 42.09% sequence identity with 18 peptides of the palustrin-2 family. Peptides that belong to the palustrin-2 family exhibit a variety of potent biological activities, including antimicrobial and anticancer activities, and therefore represent exciting candidates for novel drug development.

Palustrin-Ca is 31 amino acid residues long, of which 5 are lysine, 4 are glycine and 4 are leucine. The peptide has a cyclic disulphide-bridged heptapeptide domain at its C-terminus, which is conserved with other peptides of the palustrin-2 family.

This study employed NMR spectroscopy and molecular modelling methods to characterise the peptide’s three-dimensional structure and obtain an ensemble of model structures. To date, palustrin-Ca is the only palustrin peptide to have had its three-dimensional structure elucidated. The results established that palustrin-Ca possesses an amphipathic, *α*-helical structure between residues Ile^6^-Ala^26^ in a 50% TFE-H_2_O mixed solvent system, with the terminal segments predominantly defined as turns, with significant coil character as well.

A molecular dynamics simulation was conducted, whereby the medoid NMR-determined structure was simulated with an SDS micelle. The simulation results suggest that palustrin-Ca preferentially adopts a position parallel to the micelle’s surface. This configuration is most energetically favourable as the peptide’s hydrophobic residues can penetrate to the micelle’s hydrophobic core, while the hydrophilic residues remain in contact with the aqueous solvent. The RDF plot in Figure 8 illustrates this clearly; the greatest values for *g*(*r*) at small distances are observed for the hydrophobic Leu, Ile, Ala and Phe residues, demonstrating the importance of hydrophobicity for a peptide’s ability to bind the target membrane. The observed preference for a position parallel to the micelle surface indicates that the peptide most probably exerts its antimicrobial activity through a non-pore-forming mechanism of action, such as the carpet model or the interfacial activity model.

This study’s results will facilitate future work focused on investigating HDP structures, and their interactions with zwitterionic lipid bilayers, in order to further our understanding of the relationships between HDP structure and function. This study has extended earlier work from our research group that led to the elucidation of a number of host-defence peptide structures, and also complements related bioinformatic studies on peptide structure and function^[51–54]^.

## Acknowledgements

CH is grateful to John O’Brien and Manuel Ruether at Trinity College Dublin for NMR facilities, University College Dublin for a Research Scholarship to PBT, ICHEC for access to supercomputer facilities. The solution structure of palustrin-Ca was deposited to the PDB at the RSCB with deposition code 7P4X.

